# *Ap2s1* mutation in mice causes familial hypocalciuric hypercalcemia type 3

**DOI:** 10.1101/2020.08.10.244244

**Authors:** Fadil M. Hannan, Mark Stevenson, Asha L. Bayliss, Victoria J. Stokes, Michelle Stewart, Kreepa G. Kooblall, Caroline M. Gorvin, Gemma Codner, Lydia Teboul, Sara Wells, Rajesh V. Thakker

## Abstract

Mutations of the adaptor protein-2 sigma subunit (*AP2S1*) gene which encodes AP2σ2, a component of the ubiquitous AP2 heterotetrameric complex involved in endosomal trafficking of the calcium-sensing receptor (CaSR), cause familial hypocalciuric hypercalcemia type 3 (FHH3). FHH3 patients have heterozygous *AP2S1* missense Arg15 mutations (p.Arg15Cys, p.Arg15His or p.Arg15Leu) with marked hypercalcemia and occasional hypophosphatemia and osteomalacia. To further characterise the phenotypic spectrum and calcitropic pathophysiology of FHH3, we used CRISPR/Cas9 genome editing to generate mice harboring the *AP2S1* p.Arg15Leu mutation, which causes the most severe FHH3 phenotype. Heterozygous (*Ap2s1^+/L15^*) mice were viable, and had marked hypercalcemia, hypermagnesemia, hypophosphatemia, and increased plasma concentrations of parathyroid hormone, fibroblast growth factor 23 and alkaline phosphatase activity, but normal pro-collagen type 1 N-terminal pro-peptide and 1,25 dihydroxyvitamin D. Homozygous (*Ap2s1^L15/L15^*) mice invariably died perinatally. The *AP2S1* p.Arg15Leu mutation impaired protein-protein interactions between AP2σ2 and the other AP2 subunits, and the CaSR. Cinacalcet, a CaSR allosteric activator, ameliorated the hypercalcemia and elevated PTH concentrations, but not the diminished AP2σ2-CaSR interaction. Thus, our studies have established a mouse model with a germline loss-of-function *AP2S1* mutation that is representative for FHH3 in humans, and demonstrated that cinacalcet corrects the abnormalities of plasma calcium and PTH.

## Introduction

Familial hypocalciuric hypercalcemia (FHH) is an autosomal dominant disorder of extracellular calcium (Ca^2+^o) metabolism characterized by lifelong increases of serum calcium concentrations, mild hypermagnesemia, normal or elevated circulating parathyroid hormone (PTH) concentrations, and inappropriately low urinary calcium excretion (urine calcium to creatinine clearance ratio (CCCR) <0.01) (1). FHH is a genetically heterogeneous disorder comprising three reported variants. FHH types 1 and 2 (FHH1, OMIM #145980; FHH2, OMIM #145981) are generally associated with mild asymptomatic hypercalcemia and caused by loss-of-function mutations of the calcium-sensing receptor (CaSR), a G-protein coupled receptor (GPCR), and G-protein subunit α11 (Gα_11_), respectively, which are pivotal for regulating PTH secretion and renal tubular calcium reabsorption (2). In contrast, FHH type 3 (FHH3, OMIM #600740) is associated with a more severe biochemical phenotype that is characterized by significantly higher serum calcium and magnesium concentrations and a significantly reduced CCCR, when compared to FHH1 (3). Furthermore, FHH3 may be associated with symptomatic hypercalcemia and reduced bone mineral density (BMD), and occasionally also osteomalacia, short stature, congenital heart defects and neurodevelopmental disorders (3–6). FHH3 is caused by germline heterozygous loss-of-function mutations of the *AP2S1* gene, which is located on chromosome 19q13.3 and encodes the AP2σ2 protein (7). *AP2S1* mutations have been reported in ~70 FHH probands to-date, and affected individuals harbor a mutation affecting the AP2σ2 Arg15 residue, which may give rise to an p.Arg15Cys, p.Arg15His or p.Arg15Leu missense mutation (3, 5–11). FHH3 patients harboring the p.Arg15Leu AP2σ2 mutation have been reported to have greater hypercalcemia and to present at an earlier age than probands with p.Arg15Cys or p.Arg15His AP2σ2 mutations (3).

The AP2σ2 protein is evolutionarily highly conserved (7), and forms part of the ubiquitously expressed heterotetrameric adaptor protein-2 (AP2) complex, which also comprises AP2α, AP2β2 and AP2μ2 subunits (12). The AP2 complex plays a pivotal role in clathrin-mediated endocytosis by initiating the formation of clathrin-coated vesicles, which leads to trafficking of plasma membrane constituents to endosomes (13, 14). AP2σ2 contributes to the AP2 core structure (15), which binds to transmembrane cargo proteins such as GPCRs. Consistent with this, AP2σ2 has been shown to regulate CaSR endocytosis, and the FHH3-causing p.Arg15Cys, p.Arg15His and p.Arg15Leu mutations have all been demonstrated to impair CaSR endocytosis, thereby decreasing signalling from the endosomal CaSR (16).

We have sought to establish a mouse model to: facilitate investigation of the *in vivo* roles of the AP2σ2 protein; further characterise the phenotypic spectrum and calcitropic pathophysiology of FHH3; and evaluate CaSR-targeted therapy for this disorder. Mice harboring the AP2σ2 p.Arg15Leu mutation were generated, as this is associated with the clinically most severe phenotype in FHH3 patients.

## Results

### Generation of mice harboring an Ap2s1 mutation, p.Arg15Leu

Mutant mice on a C57BL/6J strain background were generated using CRISPR/Cas9 genome editing, as reported (17). Founder mice harbored a G-to-T transversion at c.44 within exon 2 of the *Ap2s1* gene, which was predicted to lead to a missense substitution of Arg, encoded by CGC, to Leu, encoded by CTC, at *Ap2s1* codon 15 (Figure 1). F1 generation mice were shown to harbor WT (Arg15) and mutant (Leu15) *Ap2s1* alleles, and mice derived from intercrosses of heterozygous mutant mice showed the expected Mendelian inheritance ratio of 1:2:1 at birth for the WT (*Ap2s1*^+/+^), heterozygous (*Ap2s1 ^+/L15^*), and homozygous (*Ap2s1^L15/L15^*) genotypes, respectively, which were confirmed by DNA sequence analysis (Table 1, Figure 1). WT and *Ap2s1^+/L15^* mice were viable and survived into adulthood (Table 1). However, >85% of *Ap2s1^L15/L15^* mice did not survive into adulthood (Table 1), and most died within 48 hours after birth. Because of the high rate of homozygote neonatal lethality, WT and *Ap2s1^+/L15^* mice were generated for subsequent studies by backcrossing *Ap2s1^+/L15^* mice onto the WT C57BL/6J strain background.

**Figure 1.**
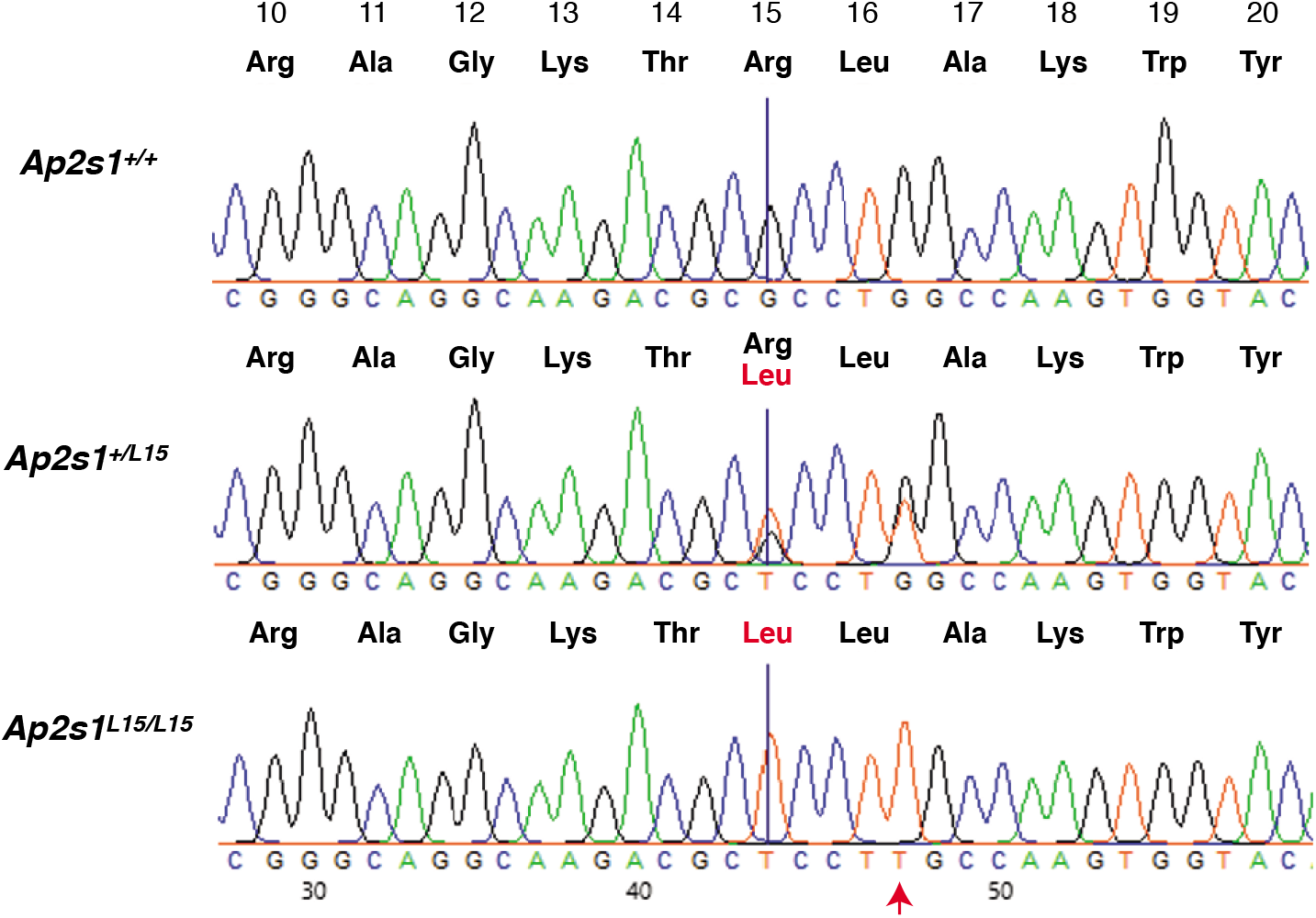
DNA sequence analysis of the CRISPR/Cas9 engineered *Ap2s1* mutation, p,Arg15Leu. The engineered G-to-T transversion at c.44 within exon 2 of *Ap2s1* is indicated by a blue line. The DNA sequence chromatograms show that WT (*Ap2s1^+/+^*) mice are homozygous G/G, the heterozygous mutant *Ap2s1^+/L15^* mice are G/T, and the homozygous mutant *Ap2s1^L15/L15^* mice are T/T. The c.G44T transversion is predicted to lead to a missense substitution of Arg (black), encoded by CGC, to Leu (red), encoded by CTC, at *Ap2s1* codon 15. A synonymous substitution, c.G48T (red arrow), was also engineered into the mutant *Ap2s1* allele, to protect the engineered allele from further re-processing by CRISPR/Cas9 reagents.

**Table 1.** Proportion of offspring bred from crosses of *Ap2s1^+/L15^* x *Ap2s1^+/L15^* mice.

| Genotype | Number of offspring<br>observed at birth<br>(n=72) | Number of offspring<br>observed at weaning<br>(~3 weeks old) (n=48) | Number of offspring observed<br>at adulthood (>10 weeks old)<br>(n=46) |
| --- | --- | --- | --- |
| +/+ (WT) | 17 (24%) | 14 (29%) | 14 (31%) |
| +/*L15* (Het) | 38 (52%) | 30 (63%) | 30 (65%) |
| *L15/L15* (Hom) | 17 (24%) | 4 (8%)** | 2 (4%***) |

WT, wild type; Het, Heterozygotes; Hom, Homozygotes. Heterozygous crosses lead to the expected Mendelian inheritance ratio of 1:2:1 for WT: Het: Hom offspring at birth. However, the observed ratio of offspring genotypes at weaning and adulthood was significantly different from the expected ratio, due to homozygote mice having reduced viability beyond the perinatal period. \*\*p<0.01 and \*\*\* p<0.001 (by binomial distribution analysis).

### Phenotype of mice harboring the Ap2s1 mutation, p.Arg15Leu

Adult *Ap2s1^+/L15^* mice, aged 12-22 weeks, showed no gross morphological abnormalities, although male *Ap2s1^+/L15^* mice had a significantly reduced body weight when compared to age-matched WT male litter-mates, whereas female *Ap2s1^+/L15^* mice had a normal body weight (Table 2). Activities such as eating, drinking, grooming, moving and interacting with cage-mates were observed to be similar between *Ap2s1^+/L15^* mice and their WT littermates. Plasma biochemical analysis showed that male and female *Ap2s1^+/L15^* mice had marked hypercalcemia in association with increased PTH concentrations and hypophosphatemia (Table 2, Figure 2A-C). These findings were accompanied by significant increases in plasma magnesium and alkaline phosphatase (ALP) activity (Table 2, Figure 2D-E). Male and female *Ap2s1^+/L15^* mice also showed marked increases in plasma fibroblast growth factor 23 (FGF23), which were not associated with alterations in plasma 1,25-dihydroxyvitamin D concentrations (Table 2, Figure 2F). Urine biochemical analysis showed that female *Ap2s1^+/L15^* mice had significantly reduced 24 hour urine calcium excretion, whereas male *Ap2s1^+/L15^* mice showed a significantly increased fractional excretion of phosphate, but no alterations of urine calcium excretion when compared to WT mice (Table 2, Figure 2G-H). Bone metabolism was assessed by whole body dual-energy X-ray absorptiometry (DXA), and by measurement of the pro-collagen type 1 N-terminal pro-peptide (P1NP) bone turnover marker, as reported (18, 19). Bone mineral content (BMC) corrected for body weight, BMD, and bone turnover in male and female *Ap2s1^+/L15^* mice were not significantly different to those observed in sex-matched WT mice (Table 2 and Supplemental Table 1).

**Figure 2.**
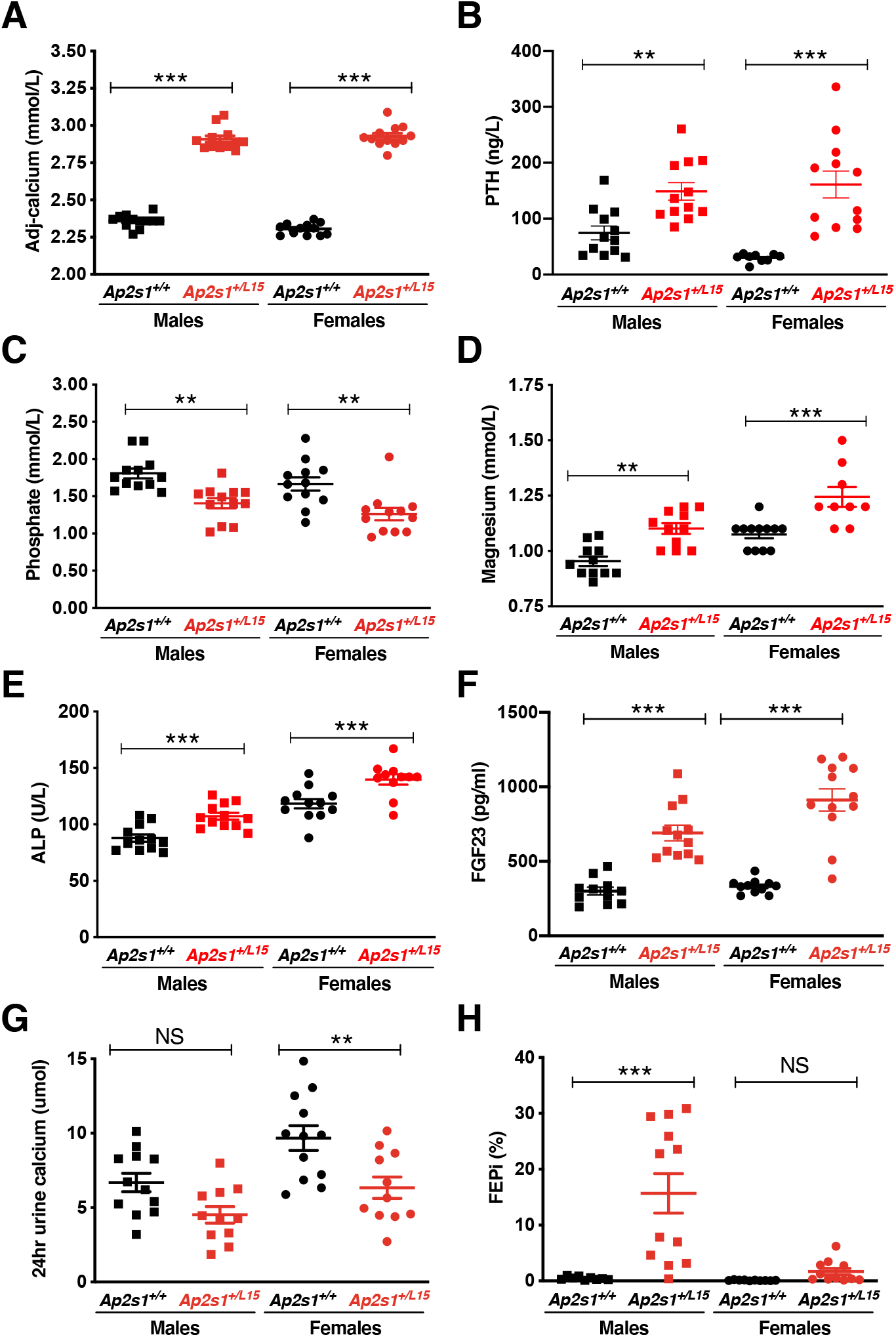
Calcitropic phenotype of adult WT (*Ap2s1^+/+^*) and *Ap2s1^+/L15^* mice, aged 15-17 weeks. (A-F) Plasma calcitropic phenotype: (A) adjusted-calcium; (B) parathyroid hormone (PTH); (C) phosphate; (D) magnesium; (E) alkaline phosphatase (ALP); and (F) fibroblast growth factor 23 (FGF23). (G-H) Urinary calcitropic phenotype: (G) 24 hour urine calcium; (H) fractional excretion of phosphate (FEPi). Mean±SEM values for respective groups (n=7-12 mice per group) are indicated. NS, non-significant; **p<0.01; ***p<0.001 for *Ap2s1^+/L15^* mice versus respective WT mice. One-way ANOVA followed by Sidak’s test for pairwise multiple comparisons were used for all analyses.

**Table 2.**
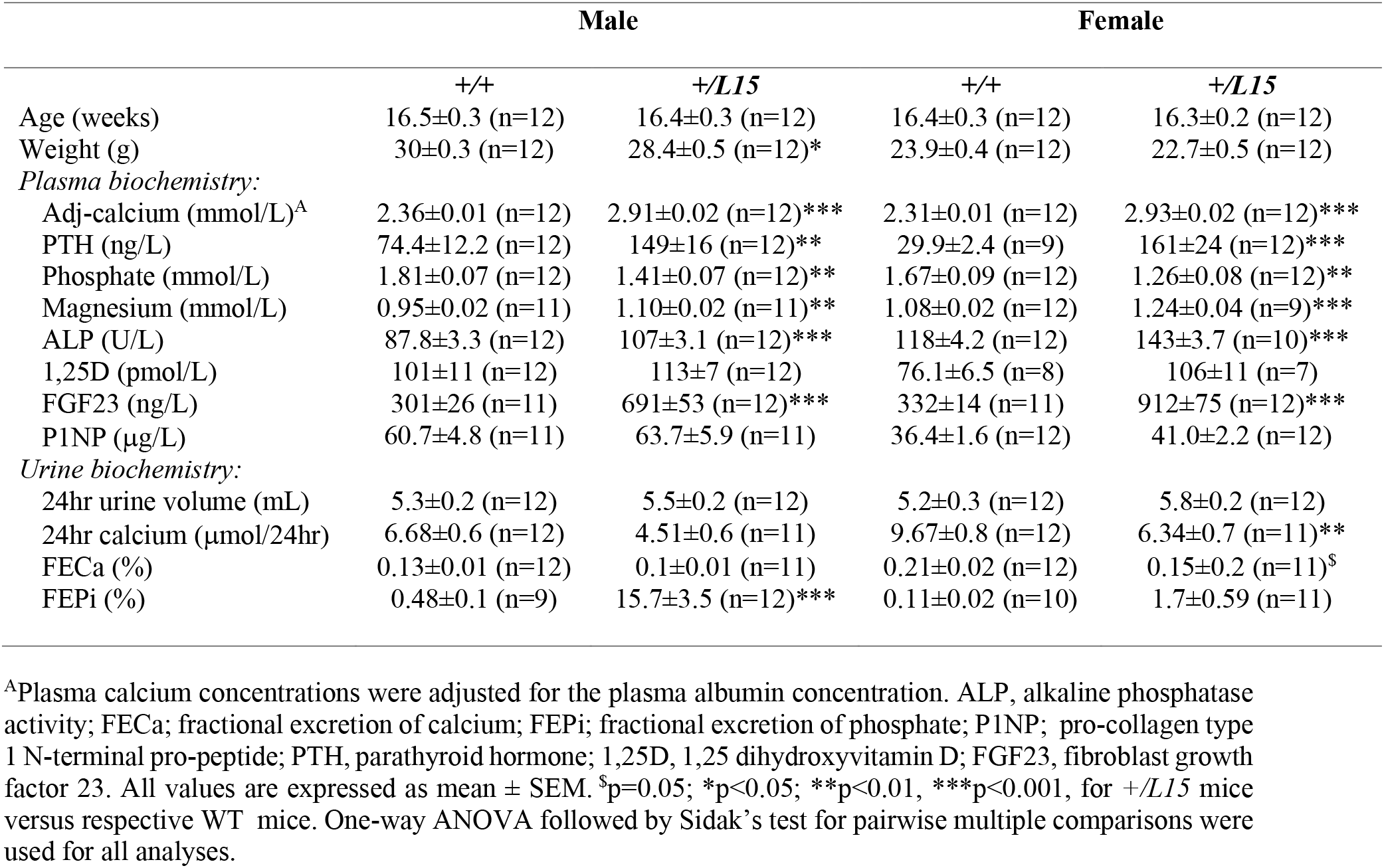
Age, body weight, and calcitropic biochemical parameters of adult WT (*+/+*) and *Ap2s1^+/L15^ (+/L15*) mice, aged 15-17 weeks.

FHH3 patients have been reported to have age-related increases in PTH concentrations (4), and we therefore assessed plasma PTH concentrations in young WT and *Ap2s1^+/L15^* mice aged 8 weeks, and also when they were mature adult mice aged 16 weeks. This analysis demonstrated an age-related increase in plasma PTH for female *Ap2s1^+/L15^* mice, which was not observed in female WT mice, or in male WT or mutant mice (Supplemental Table 2). This age-related increase in plasma PTH in female *Ap2s1^+/L15^* mice, was not associated with alterations in plasma calcium or phosphate concentrations (Supplemental Table 2).

Non-calcitropic abnormalities were observed in female *Ap2s1^+/L15^* mice, only. Thus, female *Ap2s1^+/L15^* mice had significant decreases in plasma urea and cholesterol, and a significant increase in plasma bilirubin (Supplemental Table 3). These alterations in non-calcitropic biochemical parameters were not observed in male *Ap2s1^+/L15^* mice (Supplemental Table 3).

Two male homozygous (*Ap2s1^L51/L15^*) mice survived into adulthood, and these were found to have plasma calcium concentrations >10 SD and >5 SD above the mean values of age-matched male WT and *Ap2s1^+/L15^* mice, respectively (Supplemental Table 4).

### Effect of cinacalcet on the hypercalcemia of mice harboring the Ap2s1^+/L15^ mutation

Cinacalcet, which is a CaSR positive allosteric modulator, and also known as a calcimimetic (20), has been reported to rectify impaired CaSR signalling due to FHH3-causing *AP2S1* mutations (10), and to decrease serum calcium concentrations in three FHH3 patients harboring the *AP2S1* Arg15Leu mutation (10, 21, 22). To ascertain the dose-dependent effects as well as the immediate and later actions of cinacalcet on the hypercalcemia of FHH3, we administered single oral bolus doses of 0, 30, 60 and 120 mg/kg cinacalcet to *Ap2s1^+/L15^* mice, and measured plasma PTH and calcium at 30 and 120 min post-dose. All cinacalcet doses significantly decreased plasma concentrations of PTH and calcium compared to mice given drug vehicle alone (Figure 3), and dose-dependent effects were not observed. We next treated WT and *Ap2s1^+/L15^* mice with a single 60 mg/kg cinacalcet bolus and monitored the effects on plasma calcium, phosphate, and PTH at 0, 1, 2 and 4 hours post-dose (Figure 4). Cinacalcet caused significant decreases in plasma calcium in WT and *Ap2s1^+/L15^* mice at 1 hour post-dose, and a further reduction in calcium was observed at 2 and 4 hours post-dose (Figure 4A-B). Cinacalcet treatment caused WT mice to become hyperphosphatemic, but such alterations in plasma phosphate were not observed in *Ap2s1^+/L15^* mice (Figure 4C-D). The 60 mg/kg cinacalcet dose did not significantly alter PTH concentrations in WT mice, but transiently decreased plasma PTH in *Ap2s1^+/L15^* mice at 1 hour post-dose (Figure 4E-F).

**Figure 3.**
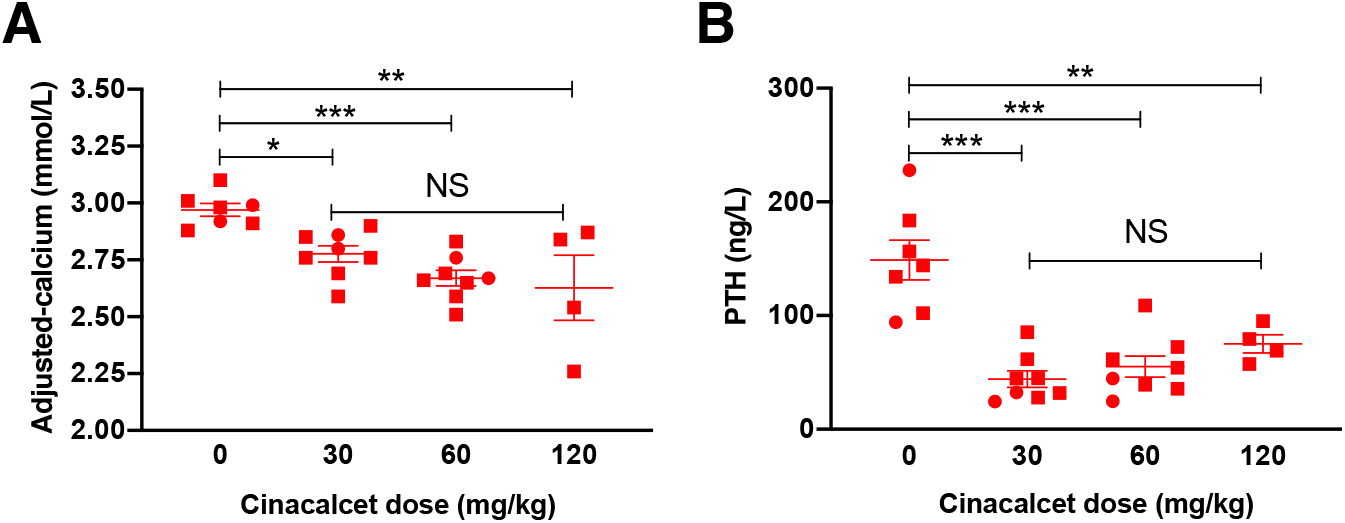
Cinacalcet dose-ranging study in adult *Ap2s1^+/L15^* mice aged 14-18 weeks. Effect of 0, 30, 60 and 120 mg/kg doses of cinacalcet on: (A) plasma albumin-adjusted calcium; and (B) plasma parathyroid hormone (PTH). Mean±SEM values for respective groups (n=4-8 *Ap2s1^+/L15^* mice per group) are indicated. Squares, males; circles, females. NS, non-significant; *p<0.05; **p<0.01; ***p<0.001 for cinacalcet-treated *Ap2s1^+/L15^* mice versus vehicle-treated *Ap2s1^+/L15^* mice. One-way ANOVA followed by Sidak’s test for pairwise multiple comparisons were used for all analyses.

**Figure 4.**
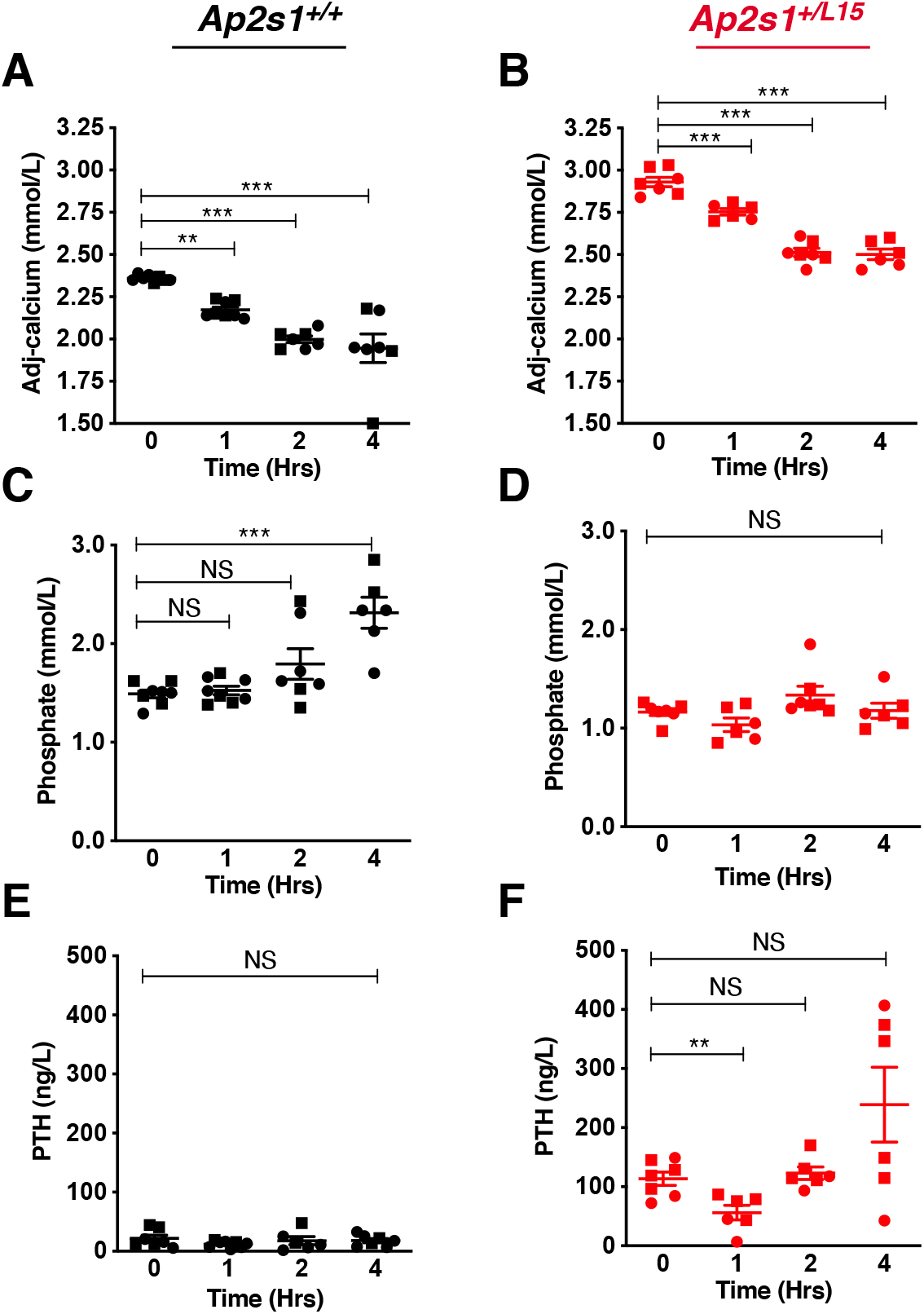
Effect of 60 mg/kg cinacalcet on plasma calcium, phosphate and parathyroid hormone at 0, 1, 2 and 4 hours post dose in adult WT (*Ap2s1^+/+^*) and *Ap2s1^+/L15^* mice, aged 15-22 weeks. (A-B) plasma albumin-adjusted calcium; (C-D) plasma phosphate; and (E-F) plasma parathyroid hormone (PTH) in *Ap2s1^+/+^* mice (black) and *Ap2s1^+/L15^* mice (red), respectively. Mean±SEM values for respective groups (n=6-8 mice per group) are indicated. Squares, males; circles, females. NS, non-significant; **p<0.01; ***p<0.001 for cinacalcet-treated mice versus respective untreated mice. Oneway ANOVA followed by Sidak’s test for pairwise multiple comparisons were used for all analyses.

### Effect of the AP2S1 p.Arg15Leu mutation on the interaction between AP2σ2 and CaSR

FHH3-causing mutations of the AP2σ2 subunit have been shown to impair CaSR endocytosis (16), and it has been postulated that this may be because of impaired interactions between AP2σ2 and the CaSR. To investigate this, we first undertook *ex vivo* co-immunoprecipitation (co-IP) analysis using renal cortical lysates from WT and *Ap2s1^+/L15^* mice (Figure 5). The CaSR was immunoprecipitated from lysates (Figure 5A) using an anti-CaSR antibody, and then probed with an anti-AP2σ2 antibody. The CaSR immunoprecipitate from WT and *Ap2s1^+/L15^* mouse kidneys showed the presence of AP2σ2, thereby confirming a protein-protein interaction between the CaSR and AP2σ2 subunit (Figure 5B). Moreover, the amount of AP2σ2 in the CaSR immunoprecipitate from *Ap2s1^+/L15^* mouse kidneys showed a >50% (p=0.057) decrease compared to kidneys from WT mice, which was suggestive of a reduced interaction between the CaSR and mutant (Leu15) AP2σ2 subunit (Figure 5C). We therefore further evaluated the effects of the FHH3-associated Arg15Leu AP2σ2 mutation on AP2σ2-CaSR interactions *in vitro,* by generating and using HEK293 cells that stably overexpressed an N-terminal FLAG-tagged CaSR (FlaC2 cells) with either the HA-tagged WT (Arg15) AP2σ2 subunit (Arg15-AP2σ2-FlaC2 cells) or HA-tagged mutant (Leu15) AP2σ2 subunit (Leu15-AP2σ2-FlaC2 cells). Co-IP analysis using anti-FLAG and anti-HA antibodies showed a significant reduction of >50% in the amount of AP2σ2 present in the FLAG-CaSR immunoprecipitate from mutant (Leu15-AP2σ2-FlaC2) cells compared to WT (Arg15-AP2σ2-FlaC2) cells (p<0.05) (Figure 6A-B). Thus, these studies demonstrate that the *AP2S1* p.Arg15Leu mutation diminishes the protein-protein interaction between CaSR and the AP2σ2 subunit. Cinacalcet improves the calcitropic phenotype of *Ap2s1^+/L15^* mice (Figure 4A-B) and FHH3 patients (10, 21, 22), and we therefore evaluated whether this calcimimetic compound can rescue the interaction between the CaSR and mutant Leu15-AP2σ2 subunit *in vitro.* WT (Arg15-AP2σ2-FlaC2) cells and mutant (Leu15-AP2σ2-FlaC2) cells were treated with 10nM cinacalcet, as this dose has been reported to rectify the signalling responses of cells expressing FHH3 mutant proteins *in vitro* (10), and the AP2σ2-CaSR interaction assessed by co-IP analysis using anti-FLAG and anti-HA antibodies. Cinacalcet treatment did not alter the amount of AP2σ2 present in the FLAG-CaSR immunoprecipitates from either WT (Arg15-AP2σ2-FlaC2) cells or mutant (Leu15-AP2σ2-FlaC2) cells (Figure 6C-D), thereby indicating that this calcimimetic does not influence the WT or mutant AP2σ2-CaSR interactions.

**Figure 5.**
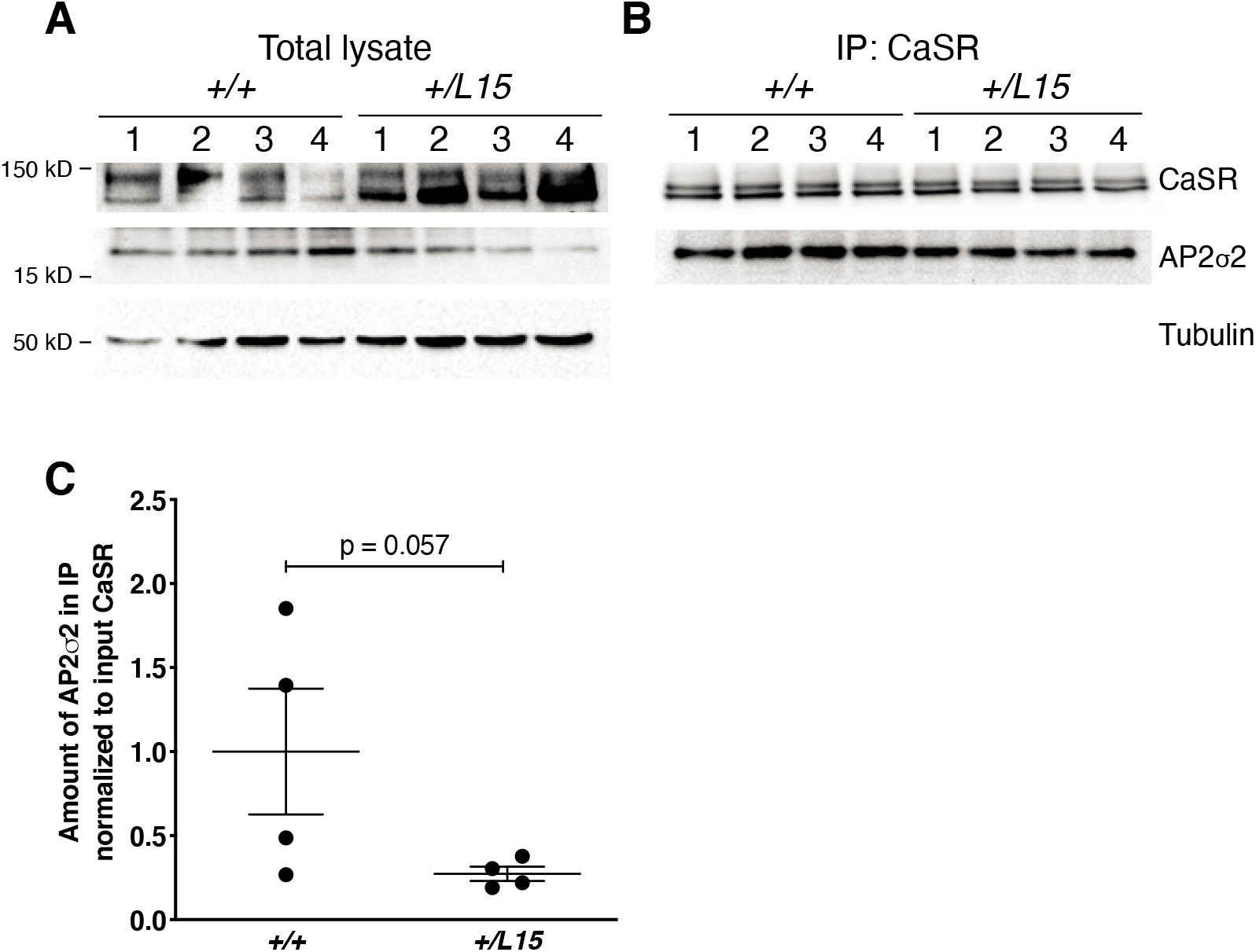
Co-immunoprecipitation analysis of the AP2σ2-CaSR interaction in *Ap2s1^+/+^* (*+/+*) and *Ap2s1^+/L15^ (+/L15*) mouse kidneys. A. Immunoprecipitation using an anti-CaSR antibody and kidney cortex lysates from n=4 *Ap2s1^+/+^* and n=4 *Ap2s1^+/L15^* female mice (numbered 1-4 in panels A-B). The amount of protein in (A) the total lysates and (B) the precipitated immune complexes (IP:CaSR) was analyzed by Western blotting using anti-CASR, anti-AP2σ2 and anti-tubulin antibodies. (C) Densitometry of Western blots to quantify AP2σ2 in the immunoprecipitate from *Ap2s1^+/+^* and *Ap2s1^+/L15^* mice normalized to the amount of CaSR in the total lysate (pre-normalized to tubulin). Mean±SEM values are indicated. Data were analyzed using a one-tailed Mann-Whitney U test.

**Figure 6.**
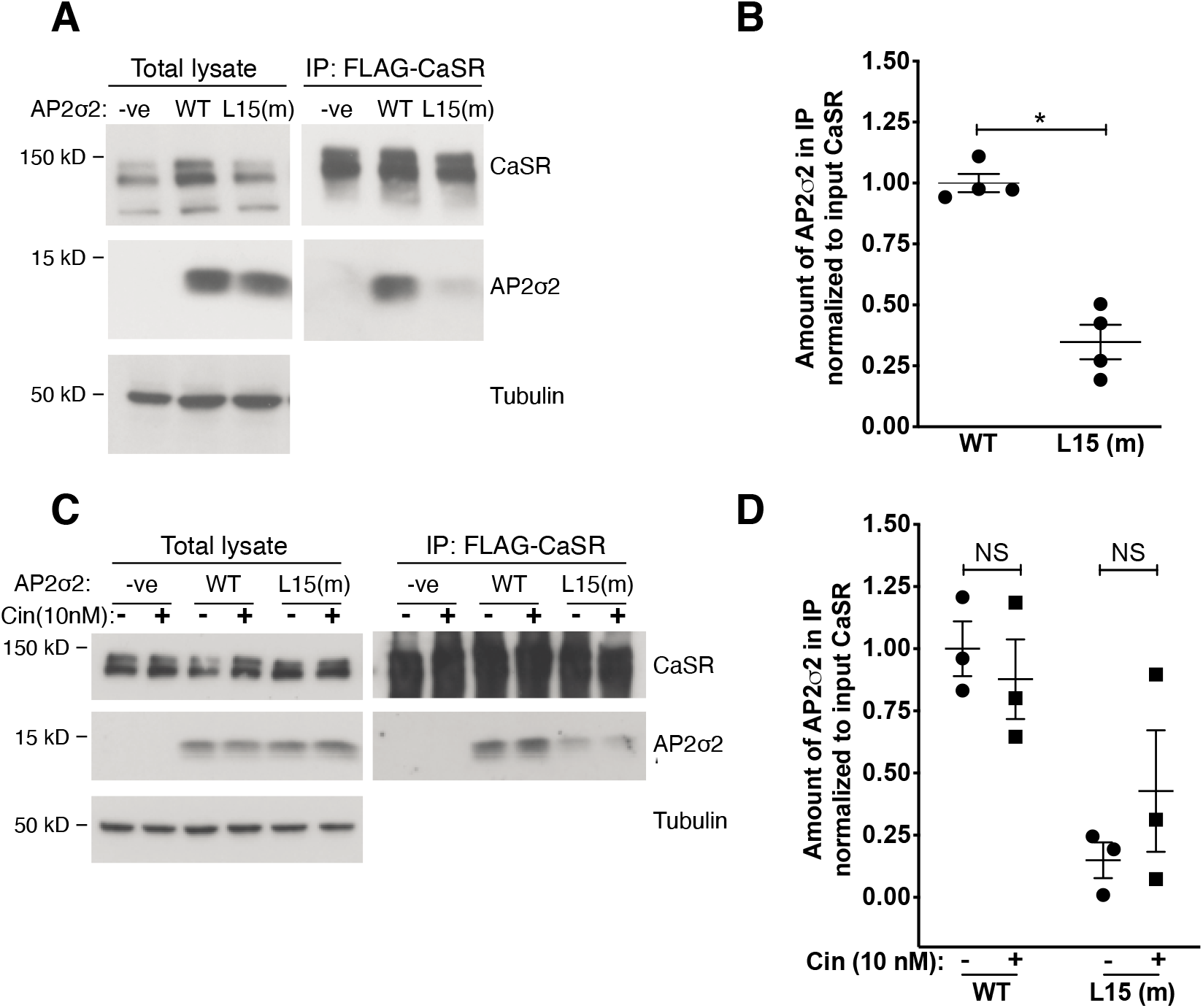
Co-immunoprecipitation analysis of the AP2σ2-CaSR interaction in HEK293 cells. (A) Immunoprecipitation using an anti-FLAG antibody and HEK293 cells stably expressing either: FLAG-tagged CaSR and HA-tagged WT AP2σ2 (WT); FLAG-tagged CaSR and HA-tagged mutant (m) Leu15 AP2σ2 (L15(m)); or FLAG-tagged CaSR alone (-ve). The amount of protein in the total lysates and precipitated immune complexes (IP:FLAG-CaSR) was analyzed by Western blotting using anti-CASR, anti-HA, and anti-tubulin antibodies. (B) Densitometry of Western blots to quantify AP2σ2 in the immunoprecipitate of WT and L15 (m) cells normalized to the amount of CaSR in the total lysate (prenormalized to tubulin). Mean±SEM values are indicated and data were analyzed using the Mann-Whitney U test. *p<0.05. (C) Immunoprecipitation using an anti-FLAG antibody in-ve, WT and L15 (m) cells treated with 10 nM cinacalcet (+) and compared to untreated cells (-). The amount of protein in the total lysates and precipitated immune complexes (IP:FLAG-CaSR) was analyzed by Western blotting using anti-CASR, anti-HA, and anti-tubulin antibodies. (D) Densitometry of AP2σ2 in the immunoprecipitate from cinacalcet-treated and untreated cells. Mean±SEM values are indicated, and data were analyzed using a using a two-way ANOVA with Bonferroni correction for multiple tests and Post Hoc analysis. NS, non-significant.

### Effect of the AP2S1 p.Arg15Leu mutation on AP2σ2 interactions with other AP2 complex subunits

The AP2σ2 subunit interacts with the AP2α, AP2ß2 and AP2μ2 subunits to form the heterotetrameric AP2 complex (23), and we therefore assessed the effects ofthe FHH3-associated Arg15Leu AP2σ2 mutation on the interactions with these other AP2 complex subunits. We undertook co-IP analysis using an anti-HA antibody and lysates from the WT (Arg15-AP2σ2-FlaC2) cells or mutant (Leu15-AP2σ2-FlaC2) cells, as these stably overexpress HA-tagged WT or mutant (Leu15) AP2σ2 proteins, and have endogenous expression of the AP2α, AP2ß2 and AP2μ2 subunits (Figure 7A). The amounts of AP2σ2 detected in the immunoprecipitate from WT (Arg15-AP2σ2-FlaC2) cells or mutant (Leu15-AP2σ2-FlaC2) cells was not significantly different (Figure 7A-B), but significant reductions of >50% in the amount of endogenous AP2α, AP2ß2 and AP2μ2 subunits were observed in the immunoprecipitate from mutant (Leu15-AP2σ2-FlaC2) cells compared to WT (Arg15-AP2σ2-FlaC2) cells (p<0.05) (Figure 7C-E). Thus, these findings indicate that the *AP2S1* FHH3-associated p.Arg15Leu mutation impairs the interaction between AP2σ2 and the other subunits of the AP2 heterotetrameric complex.

**Figure 7.**
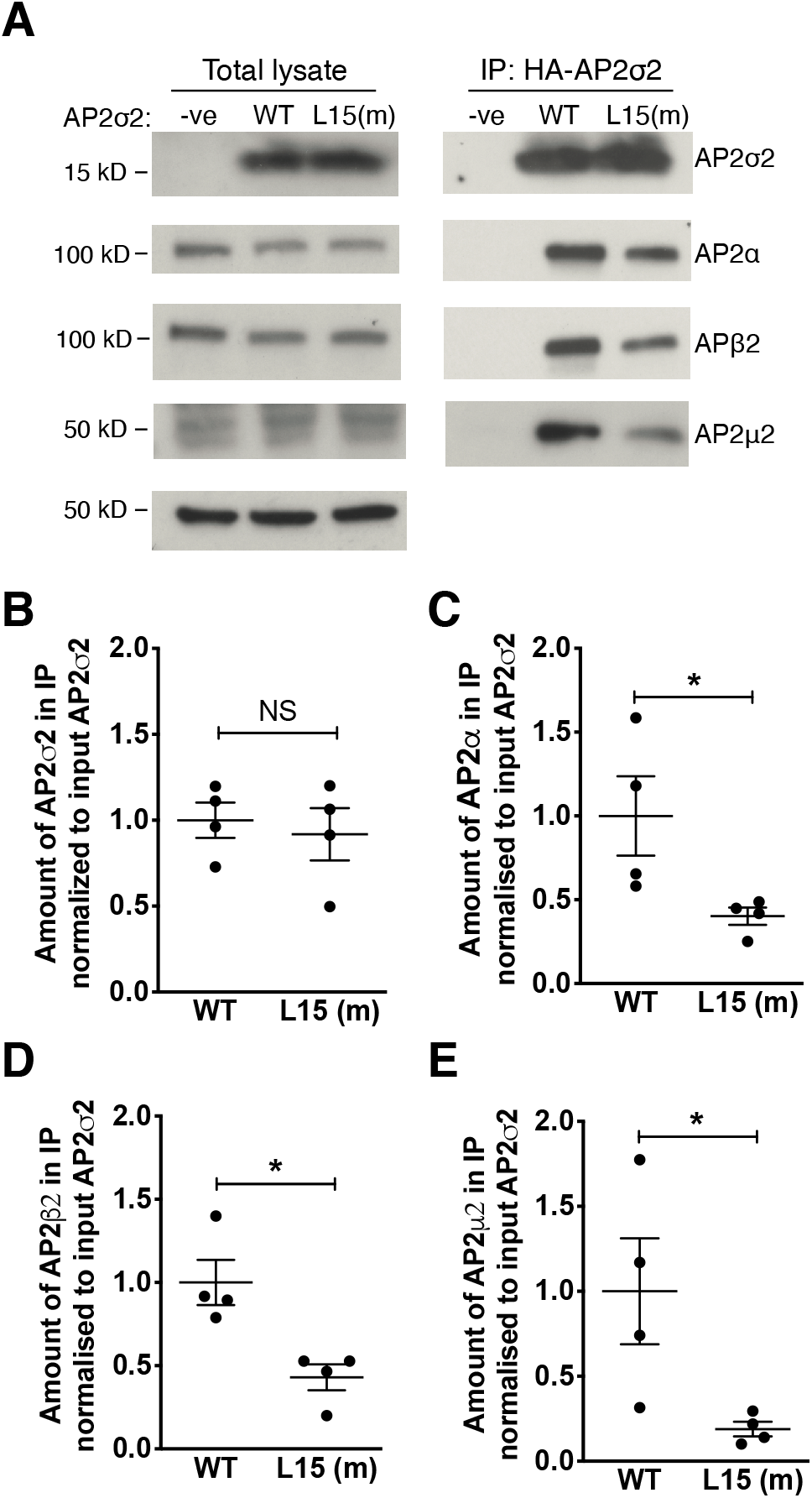
Co-immunoprecipitation analysis of the interaction between AP2σ2 and other subunits of the AP2 complex in HEK293 cells. (A) Immunoprecipitation using an anti-HA antibody and HEK293 cells stably expressing either: FLAG-tagged CaSR and HA-tagged WT AP2σ2 (WT); FLAG-tagged CaSR and HA-tagged mutant (m) Leu15 AP2σ2 (L15 (m)); or FLAG-tagged CaSR alone (-ve). The amount of protein in the total lysates and precipitated immune complexes (IP:HA-AP2σ2) was analyzed by Western blotting using anti-HA, anti-AP2α, AP2β2, AP2μ2, and anti-tubulin antibodies. (B-E) Densitometry of the Western blotting to quantify; (B) AP2σ2; (C) AP2α; (D) AP2β2; and (E) AP2μ2 in the immunoprecipitate of WT and L15 (m) cells, normalized to the amount of AP2σ2 in the total lysate (pre-normalized to tubulin). Mean±SEM values are indicated, and data were analyzed using the Mann-Whitney U test. NS, non-significant; *p<0.05.

## Discussion

We have established by the use of CRISPR/Cas9 genome editing, a mouse model for FHH3, and this will enable the calcitropic roles of AP2σ2 and endosomal trafficking of the CaSR to be further evaluated together with pursuit of pathophysiological studies that are difficult to undertake in patients with this condition. Our results revealed that *Ap2s1^+/L15^* mice, which harbored a germline heterozygous *Ap2s1* p.Arg15Leu mutation, had a similar plasma biochemical phenotype to that reported for FHH3 patients, who have heterozygous loss-of-function *AP2S1* missense Arg15 mutations (p.Arg15Cys, p.Arg15His or p.Arg15Leu), and with those having the AP2σ2 p.Arg15Leu mutation being affected with the severest hypercalcemia (3, 7). Thus, *Ap2s1^+/L15^* mice had substantial hypercalcemia with mean plasma calcium concentrations that were >0.5 mmol/L (>20%) above that of WT littermates (Table 2, Figure 2). These findings contrast with other monogenic FHH mouse models; for example *Casr^+/−^* and *Gna11^+/−^* mutant mice, which are respective models for FHH1 and FHH2, typically have milder hypercalcemia with plasma or serum calcium concentrations that are <10% above that of the WT values (19, 24). The *Ap2s1^+/L15^* mice were also hypermagnesemic, which is consistent with the phenotype of FHH3 patients (3). In addition, *Ap2s1^+/L15^* mice had significantly increased plasma PTH concentrations in association with hypophosphatemia (Table 2, Figure 2), and this is in agreement with the findings from studies of two large FHH3 kindreds from Oklahoma (FHH_Ok_) and Northern Ireland (FHH_NI_), which have demonstrated that hypercalcemic family members, compared to normocalcemic relatives, have significantly increased serum PTH concentrations with mild hypophosphatemia (4, 25). Moreover, affected males and females from the FHH_Ok_ kindred were reported to have an age-related increase in PTH (4), and such an age-related increase in PTH was also observed in the female, but not male, *Ap2s1^+/L15^* mice (Supplementary Table 2). The etiology of this age-related increase in PTH, which has not been reported in FHH1 and FHH2 kindreds, and basis of the gender differences remain to be elucidated. However, the increased plasma FGF23 concentrations that were observed in the *Ap2s1^+/L15^* mice (Table 2, Figure 2) are likely to have a role in the etiology of hypophosphatemia. Thus, the increased plasma FGF23 concentrations, which are likely the result of elevations in PTH that promote osteoblast production of FGF23 (26), will act on the kidneys to increase excretion of phosphate that will lead to hypophosphatemia. Interestingly, patients with primary hyperparathyroidism are reported to have increased plasma FGF23 concentrations (27), although these have not been assessed in FHH patients to-date, and thus our findings from the *Ap2s1^+/L15^* mice indicate that such measurements of FGF23 are warranted in FHH patients.

The calcitropic phenotype of *Ap2s1^+/L15^* mice showed some differences to FHH3 patients. Thus, only female *Ap2s1^+/L15^* mice were hypocalciuric, whereas male and female FHH3 patients have been reported to be hypocalciuric (urine calcium creatinine clearance ratio <0.01) (3, 4). In addition, *Ap2s1^+/L15^* mice showed no alterations in bone turnover or whole body BMD, whereas lumbar spine and/or femoral neck BMD has been reported to be decreased in ≥50% of FHH3 patients (3, 9). Moreover, *Ap2s1^+/L15^* mice had a significant increase in ALP activity (Table 2, Figure 2), which has not been observed in FHH3 patients (3). However, the increased circulating ALP activity in the *Ap2s1^+/L15^* mice was associated with normal plasma concentrations of P1NP (Table 2), which is a bone formation marker, thereby indicating that the raised ALP of *Ap2s1^+/L15^* mice may be of extra-skeletal origin. The raised ALP *of Ap2s1^+/L15^* mice is unlikely to be of hepatic origin, as mice, in contrast to humans, express little or no ALP in the liver (28), and the increased ALP of *Ap2s1^+/L15^* mice may therefore possibly arise from a non-hepatic source such as the intestine.

AP2σ2 forms part of the heterotetrameric AP2 complex, which plays a pivotal role in clathrin-mediated endocytosis, and crystallographic studies have indicated that the WT Arg15 AP2σ2 residue is involved in binding to peptide sequences on membrane-associated cargo proteins, which contain acidic dileucine motifs such as that predicted to occur in the distal portion of the CaSR intracellular domain (15). We have previously proposed that substitution of the polar Arg15 residue with the FHH3-associated non-polar Leu15 residue would disrupt the interaction between AP2σ2 and this endocytic recognition motif of the plasma membrane-bound CaSR (7), and our co-IP studies using *Ap2s1^+/L15^* mouse kidneys and HEK293 cells stably overexpressing the AP2σ2 subunit, now provide the evidence for this specific AP2σ2-CaSR interaction and its impairment by the FHH3-associated mutant Leu15 AP2σ2 protein (Figures 5 and 6). Moreover, our co-IP studies demonstrate that the FHH3-associated mutant Leu15 AP2σ2 also impairs the interactions between AP2σ2 and the other subunits (AP2α, AP2ß2 and AP2μ2) of the AP2 heterotetramer (Figure 7), thereby highlighting the pivotal role of this mutation in disrupting clathrin-mediated endocytosis of cell-surface proteins. These findings suggest that *Ap2s1^+/L15^* mice (and FHH3 patients) may therefore exhibit non-calcitropic phenotypes in addition to the calcitropic abnormalities described above.

Indeed, our investigations of the *Ap2s1^+/L15^* mice have revealed non-calcitropic biochemical features (Supplemental Table 3). Most notably, *Ap2s1^+/L15^* females had significant reductions in both plasma total cholesterol concentration and HDL cholesterol, which is a major cholesterol fraction in mice (29). HDL cholesterol concentrations are regulated by the ATP binding cassette transporter A-1 (ABCA1), which promotes efflux of cellular cholesterol and mediates the formation of HDL particles (30), and ABCA1 mutations cause familial HDL deficiency (31). Cellular cholesterol efflux mediated by ABCA1 is influenced by clathrin-dependent and independent endocytic pathways (32), and this highlights the possibility that the AP2 complex may play a role in cholesterol efflux. Thus, the *Ap2s1* p.Arg15Leu mutation may potentially induce low plasma HDL cholesterol concentrations through effects on the ABCA1 protein, and further studies are required to elucidate this mechanism and also to assess whether FHH3 patients may have alterations of plasma lipid components. In FHH3 patients, non-calcitropic features such as short stature, congenital heart defects and neurodevelopmental disorders have been reported (3, 5, 6). Neurodevelopmental disorders are reported to affect >65% of FHH3 children, who may have mild to severe learning difficulties, and also behavioural disturbances such as autism-spectrum disorder (ASD) and attention deficit hyperactivity disorder (ADHD) (3, 5, 6). Our present study, which included juvenile and adult *Ap2s1^+/L15^* mice, did not detect any gross behavioural abnormalities, but specific behavioural, neurophysiological and cognitive assessments will be required to identify any occurrence of neurodevelopmental abnormalities in the *Ap2s1^+/L15^* mice.

*Ap2s1^L15/L15^* homozygotes, in contrast to *Ap2s1^+/L15^* mice, were sub-viable (Table 1), and most died within the early neonatal period. One possible explanation for this neonatal lethality is that the *Ap2s1^L15/L15^* mice developed neonatal severe hyperparathyroidism (NSHPT), similar to that reported for *Casr^-/-^* mice (24). NSHPT is characterised by severe hypercalcemia, skeletal demineralisation and growth retardation, and *Casr^-/-^* mice with NSHPT have been reported to die within 3-30 days after birth (24). However, most *Ap2s1^L15/L15^* homozygotes died earlier, typically within 48 hours after birth, which may suggest an alternate etiology for their neonatal lethality, such as a generalised impairment of clathrin-mediated endocytosis caused by the disrupted interactions between mutant AP2σ2 and the other subunits of the AP2 heterotetrameric complex (Figure 7C-E). Consistent with this, a missense mutation of another AP2 subunit, AP2μ2, has been reported to impair clathrin-mediated endocytosis, and to be associated with epilepsy and developmental encephalopathy (33). Two male *Ap2s1^L15/L15^* mice did survive into adulthood, and both had more severe hypercalcemia than their male *Ap2s1^+/L15^* littermates (Supplemental Table 4), thereby also indicating a dosage effect of the mutant Leu15 *Ap2s1* allele on plasma calcium concentrations.

Cinacalcet decreased plasma PTH and calcium concentrations (10, 21, 22) in *Ap2s1^+/L15^* mice, such that a single dose caused a rapid and >50% decrease in plasma PTH concentrations and a substantially lowering of plasma calcium by ~0.40 mmol/L, although treated *Ap2s1^+/L15^* mice remained mildly hypercalcemic (Figures 3 and 4). However, it is likely that longer-term cinacalcet dosing would normalise plasma calcium concentrations in *Ap2s1^+/L15^* mice consistent with reports of cinacalcet-treated FHH3 patients (10, 21, 22).

In summary, we have established a mouse model for FHH3, and shown that the germline p.Arg15Leu mutation affecting the ubiquitously expressed AP2σ2 protein leads to a predominantly calcitropic phenotype, most likely by impairing interaction of AP2σ2 with the CaSR. Moreover, we have demonstrated that cinacalcet has a rapid effect in decreasing plasma PTH concentrations and in alleviating the hypercalcemia associated with FHH3.

## Methods

### Generation of Ap2s1^R15L^ mice

Mice harbouring a c.G44T transversion (p.Arg15Leu) in the *Ap2s1* gene, which encodes the AP2σ2 protein, were generated by homology-directed repair using the CRISPR/Cas9 system, as reported (17, 34). Single-guide RNAs (sgRNAs) targeting the genomic region encoding the Arg15 residue of AP2σ2 were designed (http://crispr.mit.edu/), with the sgRNA cutting nearest to the intended change taken forward (5’-3’: CCGGGCAGGCAAGACGCGCC, protospacer adjacent motif (PAM) sequence: TGG). A single stranded DNA oligo-deoxynucleotide (ssODN) donor template of 121 nt containing the c.G44T (p.Arg15Leu) point mutation together with a synonymous substitution, c.G48T (p.Leu16Leu), to protect the engineered allele from further re-processing by CRISPR/Cas9 reagents, was purchased as an Ultramer™ DNA oligonucleotide (IDT) with 4 phosphorothioate bonds at each 5’ and 3’ extremity (5’-3’: c.ACCTCCTCGATCAGCTTCTGCTTCTCGTCGTCATCGAACTGCATGTACCACTTGGCAAG GAGCGTCTTGCCTGCCCGGTTCTGGATAAGGATGAATCGGATCTAGAGCAAGCAGGGGA GGG). Cas9 mRNA (Tebu-Bio), sgRNA and ssODN were diluted and mixed in microinjection buffer (MIB; 10 mM Tris–HCl, 0.1 mM EDTA, 100 mM NaCl, pH7.5) to the working concentrations of 100 ng/μl, 50 ng/μl each and 50 ng/μl, respectively, and micro-injected into the pronucleus of C57BL/6J zygotes, which were then implanted into three recipient CD1 dams. Founder mice harboring the targeted allele were identified by obtaining ear biopsy DNA, which was then amplified using the following primers (5’-3’): AGATGAACTAAAGCCTGGGGC and TGTTCTGTACGCAACGAGCC. Amplicons were analysed by Sanger DNA sequencing and one mosaic founder mouse was mated with WT C57BL/6J mice to produce the F1 generation of heterozygous mice. The mutant allele was characterised in the F1 generation by using PCR, Sanger DNA sequencing and ddPCR copy counting using both a universal assay (Forward primer 5’-3’: TGCTTGCTCTAGATCCGATTCATC, Reverse primer 5’-3’: TCGTCGTCATCGAACTGCAT, Probe 5’-3’: CTTATCCAGAACCGGGCAGGCA) and a p.Arg15Leu mutant-specific assay (Forward primer 5’-3’: GCATGTACCACTTGGCAAGGA, Reverse primer 5’-3’: TGCTTGCTCTAGATCCGATTCATC, Probe 5’-3’: TCTTGCCTGCCCGGTTCTGGAT) to confirm that random donor integrations had not occurred. F1 heterozygous mice were then inter-crossed to generate the initial litters of WT, *Ap2s1^+/L15^* and *Ap2s1^L15/L15^* mice for assessment of viability. Subsequent generations of WT and *Ap2s1^+/L15^* mice were established by backcrossing *Ap2s1^+/L15^* mice onto the C57BL/6J strain background. All mice were kept in accordance with Home Office welfare guidance in an environment controlled for light (12 hours light and dark cycle), temperature (21 ± 2°C) and humidity (55 ± 10%) at the Medical Research Council (MRC) Harwell Centre (35). Mice had free access to water (25 ppm chlorine) and were fed *ad libitum* on a commercial diet (RM3, Special Diet Services) that contained 1.24% calcium, 0.83% phosphorus and 2948 IU/kg of vitamin D (35).

### Compounds

Cinacalcet (AMG-073 HCL) was obtained from Cambridge Bioscience (catalog no. CAY16042) and dissolved in DMSO or a 20% aqueous solution of 2-hydroxypropyl-ß-cyclodextrin (Sigma-Aldrich, catalog no. H107), respectively, prior to use in *in vitro* and *in vivo* studies (19).

### Generation of stable cell lines

A pcDNA3 construct (Invitrogen) containing a full length human *CASR* cDNA (36) with an N-terminal FLAG tag (DYKDDDDK) was used to generate HEK293 cells stably expressing the CaSR with a N-terminal FLAG tag (FlaC cells), and maintained under G418 (geneticin) selection. Eight FlaC cell lines (FlaC1-8) were generated (Supplemental Figure 1A-B). Expression of the CaSR was confirmed by Western blotting using anti-CASR (ADD, Abcam) and anti-FLAG antibodies (ab49763, Abcam) (Supplemental Figure 1A-B). SRE, NFAT and Fluo4-AM intracellular calcium mobilization assays were performed using methods previously described (37, 38) in one of the cell lines - FlaC2 cells - to confirm a response to extracellular calcium stimulation (Supplemental Figure 1C), and therefore functionality of the CaSR signalling pathway. FlaC2 cells stably expressing WT (WT(Arg15)-AP2σ2 FlaC2) or mutant (Leu15-AP2σ2-FlaC2) AP2σ2 proteins, were then generated using a pcDNA5 construct containing a full-length human *AP2S1* cDNA (16) with a C-terminal HA tag (YPYDVPDYA), and maintained under hygromycin and geneticin selection. Site directed mutagenesis using the Q5 Site-Directed Mutagenesis kit (New England Biolabs) and *AP2S1* specific primers (Thermo Fisher Scientific) were used to generate the mutant (Leu15) *AP2S1* construct. Stably transfected WT-AP2σ2 FlaC2, mutant Leu15-AP2σ2-FlaC2, and control FlaC2 cells were cultured in DMEM media supplemented with 10% FCS. For drug compound studies, the cells were treated with 10nM cinacalcet, or vehicle (DMSO), for 15 min prior to being washed with PBS and lysed for co-IP analysis.

### Co-immunoprecipitation and Western Blot Analysis

Cells were lysed in ice-cold lysis buffer (0.5% NP40, 135mM NaCl, 20mM Tris pH7.5, 1mM EDTA, 1x protease inhibitor (Sigma-Aldrich) 2mM Na3VO4, 10mM NaF) and debris removed by centrifugation. Immunoprecipitations using anti-CASR antibodies, were performed by mixing lysates with antibody for 30 min at 4°C prior to addition to protein G agarose beads (Cell Signalling Technology) and further mixing for 2 hours. Alternatively, lysates were mixed with anti-FLAG-sepharose or anti-HA-sepharose beads (Cell Signalling Technology) and mixed for 2 hours. Beads were then washed 5x with lysis buffer and proteins prepared in 4x Laemmli loading dye and resolved using 10% SDS-PAGE gel electrophoresis. Proteins were transferred to polyvinylidene difluoride membrane and probed with primary antibodies (CASR (ADD, Abcam), AP2σ2 (Ab92380, Abcam), FLAG (ab49763, Abcam), HA (CST3724, Cell Signalling Technologies), AP2α (BD610501, BD Biosciences), AP2ß2 (BD610381, BD Biosciences), AP2μ2 (Ab137727, Abcam), tubulin (Ab15246, Abcam) and calnexin (AB2301, Merck)), and HRP-conjugated secondary antibodies (715-035-150 and 711-035-152, Jackson ImmunoResearch), prior to visualisation using Pierce ECL Western blotting substrate. Tubulin or calnexin protein expression was used as a loading control. Densitometry analysis was performed by calculating the number of pixels per band using ImageJ software.

### Collection of mouse kidneys for co-immunoprecipitation studies

Kidneys were collected from *Ap2s1* and *Ap2s1^+/L15^* female mice, and snap frozen in liquid nitrogen, and subsequently stored at −80°C. The outer renal capsule was removed to access the cortex, which was dissected and lysed in ice-cold lysis buffer, as described above, for co-IP analysis.

### Plasma biochemistry and hormone analysis

Blood samples from juvenile mice (aged 8 weeks) and adult mice (aged 15-22 weeks) were collected from the lateral tail vein following application of topical local anesthesia for measurement of plasma PTH, or collected from the retro-orbital vein under isoflurane terminal anesthesia for measurement of other plasma biochemical parameters (35, 39). Plasma was separated by centrifugation at 5000 g for 10 min at 8°C, and analyzed for sodium, potassium, calcium, albumin, phosphate, magnesium, alkaline phosphatase activity, glucose, lipids, liver function tests, urea and creatinine on a Beckman Coulter AU680 analyzer, as described (39). Plasma calcium was adjusted for variations in albumin concentrations using the formula: (plasma calcium (mmol/l) – [(plasma albumin (g/l) – 30) x 0.02], as reported (35). Hormones were measured as follows: PTH using a two-site ELISA specific for mouse intact PTH (Immutopics, San Clemente, USA); 1,25-dihydroxyvitamin D by a two-step process involving purification by immunoextraction and quantification by enzyme immunoassay (Immunodiagnostic Systems); and FGF23 using a two-site ELISA kit (Kainos Laboratories), as described (19). Procollagen type 1 N-terminal propeptide (P1NP) was measured by an enzyme immunoassay (EIA) (Immunodiagnostic Systems) (18).

### Metabolic cages and urine biochemistry analysis

Mice, aged 15-17 weeks, were individually housed in metabolic cages (Techniplast), and fed *ad libitum* on water and powdered chow. Mice were allowed to acclimatise to their environment over a 72h period, as described, prior to collection of 24h urine samples (19). Urine was analyzed for calcium, phosphate, and creatinine on a Beckman Coulter AU680 analyzer (19). The fractional excretion of calcium and phosphate were calculated using the formula U_x_/P_x_*P_Cr_/U_Cr_, where U_x_ is the urinary concentration of the filtered substance (substance *x*) in mmol/L, P_x_ is the plasma concentration of substance *x* in mmol/L, U_Cr_ is the urinary concentration of creatinine in mmol/L, and P_Cr_ is the plasma concentration of creatinine in mmol/L (19).

### Skeletal imaging

BMC and BMD were measured in mice, aged 12-16 weeks, by whole body DXA scanning, which was performed on mice anesthetized by inhaled isoflurane and using a Lunar Piximus densitometer (GE Medical Systems), as reported (19). DXA images were analyzed using Piximus software (19).

### In vivo administration of cinacalcet

Mice, aged 14-22 weeks, were randomly allocated to receive cinacalcet or vehicle as a single oral gavage bolus (19). None of the mice had undergone any experimental procedures prior to dosing. Study investigators were blinded during animal handling and also when undertaking endpoint measurements. The primary experimental outcome was a change in plasma calcium at 2-hours post-dose.

### Statistical analysis

All *in vitro* studies involved n=3-4 biological replicates. Statistical analysis of *in vitro* data was undertaken using the Mann Whitney U test for two group comparisons, or a two-way ANOVA with Bonferroni correction for multiple tests and post-hoc analysis. Mouse viability was assessed by binomial distribution analysis. One-way ANOVA followed by Sidak’s test for pairwise multiple comparisons were used for all *in vivo* analyses. All analyses were performed using GraphPad Prism (GraphPad), and a value of p<0.05 was considered significant for all analyses

### Study approval

Animal studies were approved by the MRC Harwell Institute Ethical Review Committee, and were licensed under the Animal (Scientific Procedures) Act 1986, issued by the UK Government Home Office Department (PPL30/3271).

## Author Contributions

Designing research studies (F.M.H., Ma.S., A.L.B., C.M.G., G.C., L.T., S.W., R.V.T.), conducting experiments (Ma.S., A.L.B., Mi.S., G.C.), acquiring and analysing data (F.M.H., Ma.S., A.L.B., V.J.S., K.G.K., G.C.), wrote the manuscript (F.M.H., Ma.S., R.V.T.).

## Acknowledgements

This work was supported by a Wellcome Trust Investigator Award (grant number 106995/Z/15/Z) (to R.V.T.), and Wellcome Trust Clinical Training Fellowship (grant number 205011/Z/16/Z) (to V.J.S.).

