## Supplemental data for "*Ap2s1* mutation in mice causes familial hypocalciuric hypercalcemia type 3"

**Supplementary Figures and Tables**

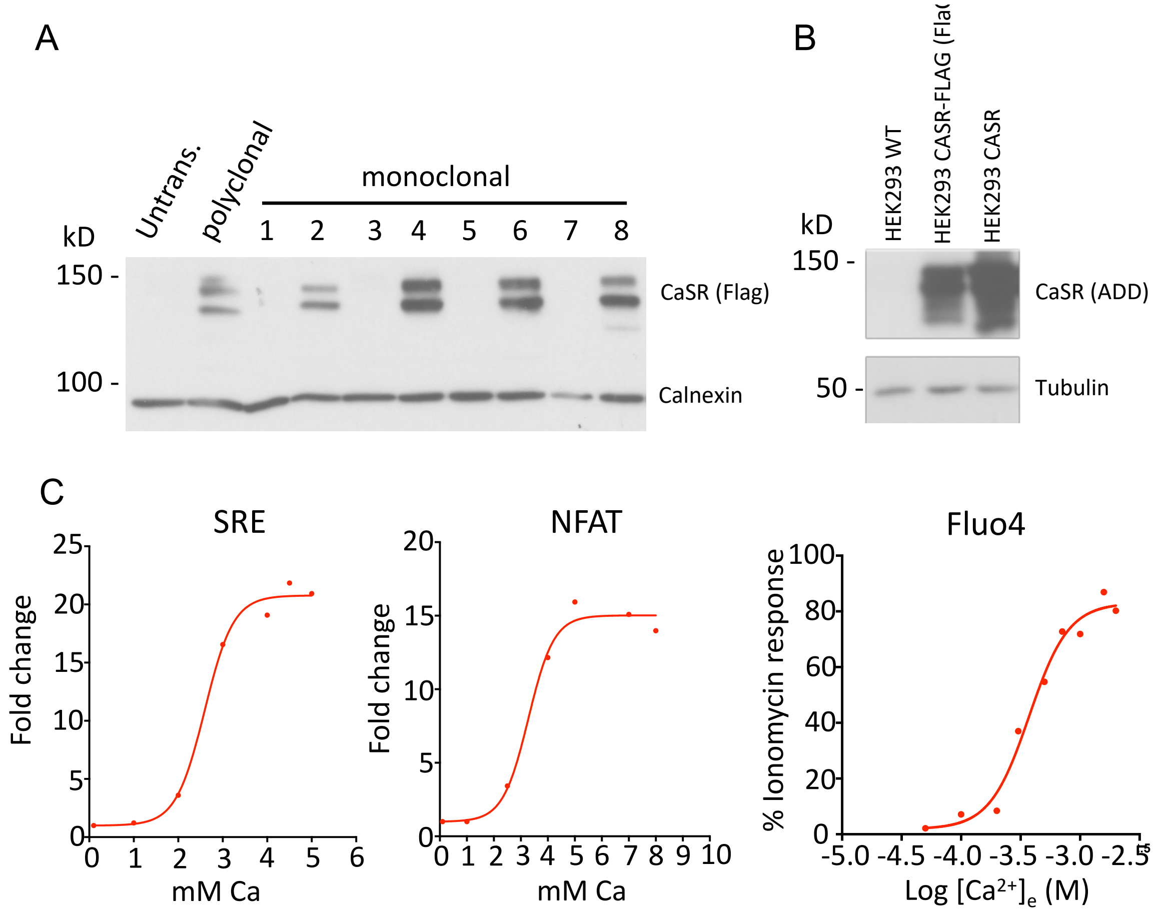


**Supplementary Figure 1.** Confirmation of functional CaSR expression in FlaC cells. (A) Western blotting analysis using an anti-FLAG antibody in 8 monoclonal stably transfected FlaC cell lines (n = 1-8) lysates, confirming CaSR expression in lines 2, 4, 6 and 8, as well as a polyclonal control line. (B) Validation of CASR expression using an anti-CASR (ADD) antibody in the FlaC2 cell line and a HEK293-CASR (without FLAG tag) control cell line is shown. (C) Functional analysis of the CaSR in FlaC2 cells using serum response element (SRE), nuclear factor of activated T cells (NFAT) and Fluo4-AM intracellular calcium mobilisation assays reveals a dose dependent response to extracellular calcium stimulation, and thus, a functional CaSR signalling pathway.

Supplementary Table 1. Whole body DXA analysis of WT *(+/+)* and *Ap2s1*$^{+/L15}$*(+/L15)* mice.

| | **Male** | | **Female** | |
|---|---|---|---|---|
| | *+/+* | ***+/L15*** | *+/+* | ***+/L15*** |
| BMC-corr (mg/g) | 23.6±0.9 (n=12) | 21.6±0.2 (n=10) | 25.4±0.4 (n=11) | 23.9±1.1 (n=11) |
| BMD (mg/cm$^2$) | 75.2±1.2 (n=12) | 71.4±0.9 (n=12) | 70.5±0.9 (n=11) | 67.6±1.6 (n=11) |
| Fat mass (%) | 15.4±1.6 (n=12) | 13.3±0.3 (n=11) | 16.4±0.9 (n=11) | 14.9±1.4 (n=11) |
| Lean mass (%) | 82.3±1.6 (n=12) | 84.9.±0.3 (n=11) | 80.2±0.9 (n=11) | 82.0±1.3 (n=11) |

BMC, bone mineral content; BMC-corr, BMC corrected for body weight; BMD, bone mineral density; DXA, dual-energy X-ray absorptiometry. All values are expressed as mean±SEM. One-way ANOVA followed by Sidak's test for pairwise multiple comparisons were used for all analyses.

Supplementary Table 2. Age-related changes in calcitropic biochemical parameters of WT *(+/+)* and *Ap2s1*$^{+/L15}$*(+/L15)* mice.

| | Male | | | | Female | | | |
|---|---|---|---|---|---|---|---|---|
| | +/+ | | *+/L15* | | +/+ | | *+/L15* | |
| | **8 weeks** | **16 weeks** | **8 weeks** | **16 weeks** | **8 weeks** | **16 weeks** | **8 weeks** | **16 weeks** |
| Adj-calcium (mmol/L)[A] | 2.49±0.05 | 2.41±0.03 | 3.11±0.01 | 2.98±0.03 | 2.48±0.04 | 2.38±0.02 | 2.94±0.03 | 2.92±0.05 |
| Phosphate (mmol/L) | 2.28±0.06 | 1.99±0.14 | 1.82±0.15 | 1.49±0.26 | 2.38±0.18 | 1.86±0.1 | 1.66±0.13 | 1.57±0.25 |
| PTH (ng/L) | 34.2±3.0 | 65.1±17 | 108±24 | 129±15 | 31.2±11.6 | 33.7±6.6 | 73.3±9.8 | 145±16** |

[A]Plasma calcium concentrations were adjusted for the plasma albumin concentration. PTH, parathyroid hormone. All values are from n=4 to 9 mice, and are expressed as mean±SEM. **p<0.01 for 8-week old mice versus respective 16-week old mice. One way ANOVA followed by Sidak's test for pairwise multiple comparisons were used for all analyses.

Supplementary Table 3. Non-calcitropic biochemical parameters of WT *(+/+)* and *Ap2s1*$^{+/L15}$ *(+/L15)* mice.

| | **Male** | | **Female** | |
|---|---|---|---|---|
| | *+/+* | ***+/L15*** | *+/+* | ***+/L15*** |
| *Electrolytes and renal:* | | | | |
| $Na^+$ (mmol/L) | 147±0.4 (n=12) | 147±0.4 (n=11) | 145±0.3 (n=12) | 144±0.5 (n=12 |
| $K^+$ (mmol/L) | 4.4±0.1 (n=12) | 4.6±0.1 (n=11) | 4.8±0.1 (n=12) | 4.9±0.1 (n=12) |
| Urea (mmol/L) | 10.7±0.4 (n=12) | 9.6±0.2 (n=12) | 10.6±0.4 (n=12) | 8.7±0.5 (n=12)** |
| Creatinine (mmol/L) | 9.8±0.3 (n=12) | 10.5±0.4 (n=12) | 10.2±0.4 (n=12) | 10.8±0.5 (n=12) |
| *Glucose and lipids*$^A$*:* | | | | |
| Glucose (mmol/L) | 13.3±0.4 (n=12) | 13.1±0.7 (n=12 | 12.1±0.7 (n=12) | 11±0.8 (n=12) |
| Total cholesterol (mmol/L) | 2.27±0.1 (n=12) | 2.17±0.04 (n=11) | 1.95±0.06 (n=12) | 1.55±0.1 (n=12)** |
| LDL-c (mmol/L) | 0.5±0.03 (n=12) | 0.4±0.01 (n=12) | 0.4±0.01(n=12) | 0.3±0.01 (n=12)$^{\$}$ |
| HDL-c (mmol/L) | 1.5±0.07 (n=12) | 1.5±0.3 (n=11) | 1.1±0.04 (n=12) | 0.8±0.08 (n=12* |
| Triglycerides (mmol/L) | 1.3±0.1 (n=12) | 1.3±0.09 (n=12) | 0.9±0.06 (n=12) | 0.9±0.06 (n=12) |
| *Liver:* | | | | |
| Total bilirubin (μmol/L) | 1.9±0.2 (n=12) | 1.5±0.1 (n=11) | 2.8±0.2 (n=12) | 3.5±0.2 (n=12)* |
| ALT (U/L) | 39.7±2.3 (n=10) | 38.2±1.7 (n=11) | 31.7±2.3 (n=12) | 30.5±1.2 (n=12) |
| AST (U/L) | 57.0±4.7 (n=12) | 57.6±5.2 (n=12) | 54.8±3.6 (n=12) | 59.8±4.4 (n=12) |

$^A$Glucose and lipid concentrations were measured in non-fasting plasma samples. ALT, alanine aminotransferase; AST, aspartate aminotransferase; HDL-c, HDL cholesterol; LDL-c, LDL cholesterol. All values are expressed as mean ± SEM. $^{\$}$p=0.05; *p<0.05; **p<0.01 for *+/L15* mice versus respective WT mice. One-way ANOVA followed by Sidak's test for pairwise multiple comparisons were used for all analyses.

Supplementary Table 4. Comparison of calcitropic parameters between male homozygous (*L15/L15*) mice (aged 18-22 weeks) and age-matched male WT (+/+) and male heterozygous (*+/L15*) mice.

| | ***Male +/+*** (n=3-4) | ***Male +/L15*** (n=5) | ***Male L15/L15*** (n=2)[A] |
|---|---|---|---|
| Adj-calcium (mmol/L)[B] | 2.36±0.01 | 2.91±0.01*** | 3.12[a,e], 3.22[a,e] |
| PTH (ng/L) | 42.0±11 | 94.3±24 | 134[b], 180[c] |
| Phosphate (mmol/L) | 1.92±0.10 | 1.32±0.05*** | 1.17[b], 1.18[b] |
| Magnesium (mmol/L) | 1.0±0.05 | 1.2±0.06 | 1.2, 1.3[d] |
| ALP (U/L) | 75.8±1.9 | 93.6±3.3* | 88[b], 150[a,e] |
| 24hr calcium (μmol/24hr) | 9.1±0.9 | 7.3±1.2 | 5.7, 8.2 |
| BMD (mg/cm$^2$) | 69.4±2.6 | 72.5±2.8 | 69.3, 69.6 |

[A]Individual values are shown for the two male homozygous mice. [B]Plasma calcium concentrations were adjusted for the plasma albumin concentration. ALP, alkaline phosphatase activity; BMD, bone mineral density; PTH, parathyroid hormone. One-way ANOVA followed by Sidak's test for pairwise multiple comparisons were used for comparisons of *+/L15* mice with WT mice. *p<0.05; ***p<0.001. Values of individual *L15/L15* mice are: [a]>10 SD; [b]>3 SD; [c]>5 SD; and [d]>2 SD away from the respective mean WT value; and [e]>5 SD away from the respective mean *+/L15* value.
